# Structure of anchorless RML prion reveals motif variation between strains

**DOI:** 10.1101/2021.12.22.473909

**Authors:** Forrest Hoyt, Heidi G. Standke, Efrosini Artikis, Cindi L. Schwartz, Bryan Hansen, Kunpeng Li, Andrew G. Hughson, Matteo Manca, Olivia R. Thomas, Gregory J. Raymond, Gerald S. Baron, Byron Caughey, Allison Kraus

**Affiliations:** Research Technologies Branch, Rocky Mountain Laboratories, National Institute of Allergy and Infectious Diseases, National Institutes of Health, Hamilton, MT 59840, USA; Department of Pathology, Case Western Reserve University School of Medicine, Cleveland, OH, USA; Laboratory of Persistent Viral Diseases, Rocky Mountain Laboratories, National Institute of Allergy and Infectious Diseases, National Institutes of Health, Hamilton, MT 59840, USA; Cleveland Center for Membrane and Structural Biology, Case Western Reserve University, Cleveland, OH, United States

**Keywords:** prion, cryo-electron microscopy, membrane, amyloid, glycan, glycophosphatidylinositol anchor, parallel in-register β-sheet, infectious, strain, species barrier

## Abstract

Little is known about the structural basis of prion strains. Here we provide a high (3.0 Å) resolution cryo-electron microscopy-based structure of brain-derived fibrils of the mouse anchorless RML scrapie strain which, like the recently determined hamster 263K strain, has a parallel in-register β-sheet-based core. However, detailed comparisons reveal that variations in shared structural motifs provide a basis for prion strain determination.

**One-sentence summary:** Cryo-electron microscopy reveals a near-atomic structure of an infectious, brain-derived murine prion fibril and strain differences.

---

Prion strains propagate as distinct infectious scrapie PrP (PrP^Sc^) conformers that can take the form of amyloid fibrils. However, the detailed structures of these conformers and how they differ between strains have been poorly understood. We recently solved a near atomic cryo-electron microscopy (cryo-EM)-based structure of a highly infectious brain-derived prion strain, i.e., 263K hamster-adapted scrapie ^1,2^. The core of this fibril has PrP monomers stacked in a parallel in-register intermolecular β-sheet (PIRIBS) based architecture. Lower resolution data (~10 Å) for the mouse anchorless RML strain (aRML, also known as anchorless Chandler) indicated a distinct fibrillar morphology, but did not allow threading of the polypeptide within the fibril core ^2^. The aRML strain provides a striking contrast to 263K. Whereas 263K PrP^Sc^ contains glycophosphatidylinositol (GPI) anchors and is heavily glycosylated, aRML PrP^Sc^ lacks GPI anchors and, consequently, are poorly glycosylated ^3,4^. Accumulation of aRML PrP^Sc^ in the brain is predominantly in extracellular amyloid plaques ^4,5^ whereas 263K is found mostly in diffuse deposits that are closely associated with cellular membranes. Although 263K fibrils have a PIRIBS architecture ^2^, aRML fibrils have been inferred by others to have a fundamentally different 4-rung β-solenoid based architecture ^6,7^.

Methodological improvements have allowed us to obtain more highly resolved cryo-EM data for aRML using single particle acquisition and helical reconstruction ^8^ with parameters given in (Methods and Supplemental Table). Fast Fourier transforms of particle 2D class averages indicated regular axial spacings of 4.9 Å, which were also visible in images of class averages (Supplemental Fig. 1a,b). The 2D class averages were used to develop 3D classifications that converged on a single core morphology. We determined a 3.0 Å resolution map of the fibril core with helical reconstruction techniques (Fig. 1b,c; Supplemental Fig. 1c & Supplemental Table). Consistent with the Fourier transforms described above, the maps indicated uniform rungs perpendicular to the fibril axis with 4.9 Å spacing (Fig. 1c,f).

**Fig. 1.**
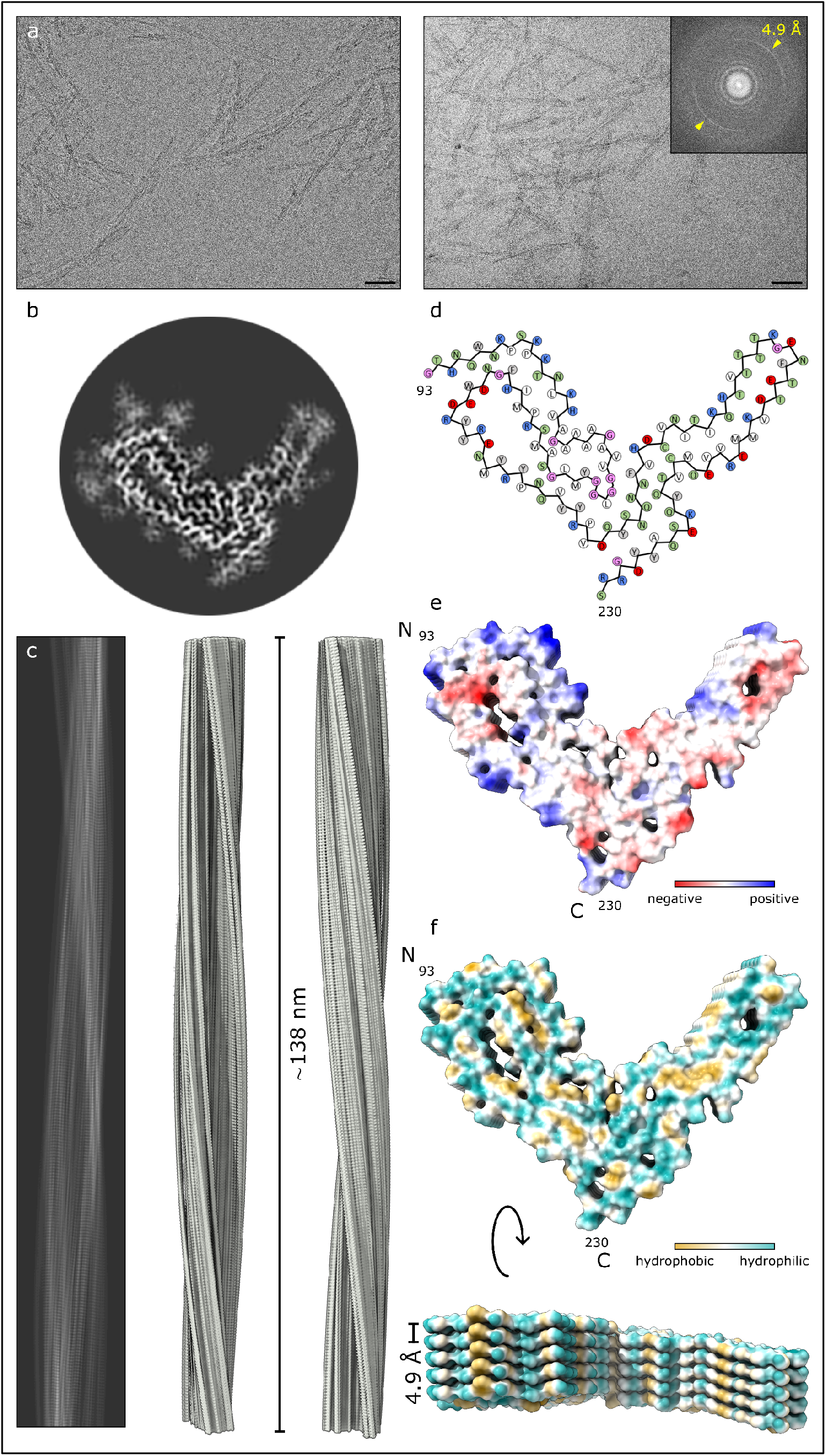
Cryo-EM-based structure of aRML fibrils. **a**, 2D cryo-EM images of aRML fibrils. Bar = 50 nm. Inset depicts associated Fourier transform showing signals from regular 4.9 Å spacings (yellow arrows). **b**, Cross-sectional view of a density map projection. **c**, Lateral view of the fibril density map with cross-over distance as indicated. **d**, Core sequence showing relative orientations of side chains. Green, polar; blue, basic; red, acidic; white, aliphatic; gray, aromatic; pink - glycine. **e**, Coulombic charge representation. **f**, Kyte-Doolittle hydrophobicity surface plot demonstrating interspersed hydrophobic interactions.

We threaded the polypeptide comprising the aRML protease-resistant core into the cryo-EM density map, with iterative real and Fourier space refinements and validation as indicated in Table 1. *In silico* energy minimization confirmed the overall stability of the structure (Methods, Supplemental Fig. 2). The resolved aRML structure comprises residues 93-230 (Fig. 1d). Some peripheral electron densities were not assigned to any of these residues (Fig. 2c, green arrows) and are assumed to reflect either tightly bound remnants of more extreme N-terminal PrP sequence after partial proteolysis, or unidentified non-PrP ligands. Peripheral densities that we previously observed adjacent to the C-terminus and N-linked glycosylation sites in the 263K prion structure ^2^ are absent in the aRML density map (Fig. 2c). As with 263K, the PrP monomers span the entire fibril cross-section and are stacked parallel and in-register perpendicular to the fibril axis with an average spacing of 4.9 Å (Fig. 1f). As detailed below, several major features of the aRML structure are analogous to, but distinct from, those in the 263K structure (Fig. 2) ^2^.

**Fig. 2.**
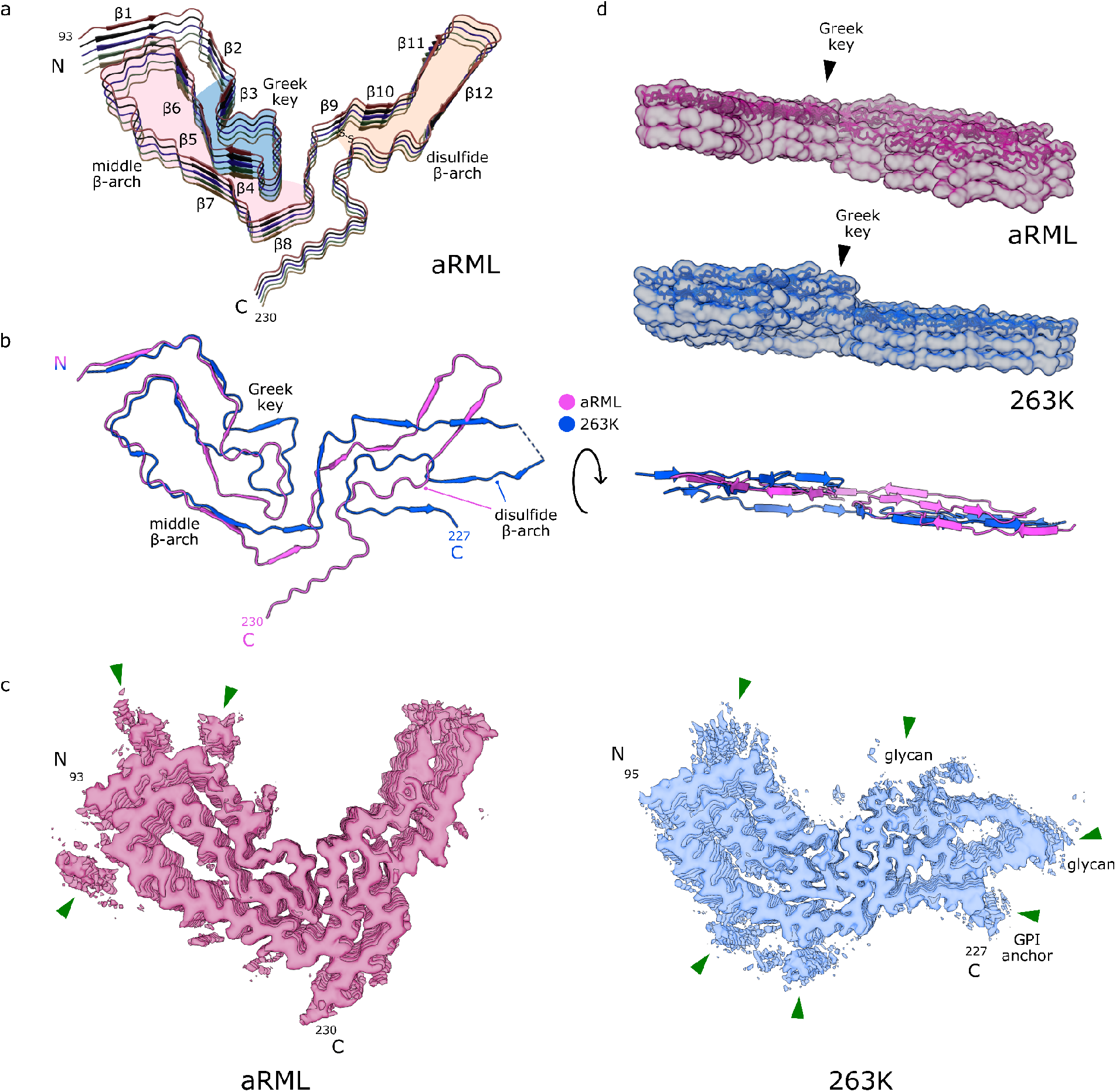
Comparison of aRML and 263K prions. **a**, Ribbon diagram of aRML core (stack of 5), with structural motifs as colored. **b**, Overlay of aRML and 263K cores. **c**, Contour EM density maps of aRML and 263K. Green arrows indicate peripheral unassigned densities associated with cationic residues in the N-terminal lobes of both strains, but absent in the aRML C-terminal lobe, consistent with aRML’s lack of glycans and glycolipid anchors. **d**, Lateral views of aRML and 263K EM maps (stacks of 3) with stick models embedded in top rung. Below is a ribbon overlay of aRML and 263K monomers indicating differences in planarity.

Residues 95-104 near the N-terminus of the aRML core sequence are held tightly against residues 140-144 by a tight interdigitation of alternate sidechains across the interface (Figs. 2c & 3a). This steric zipper-like interface is similar to that seen for analogous residues in 263K ^2^. In both aRML and 263K, the backbones of these sequences are not precisely coplanar within a given monomer but are staggered slightly with respect to one another axially.

**Fig. 3.**
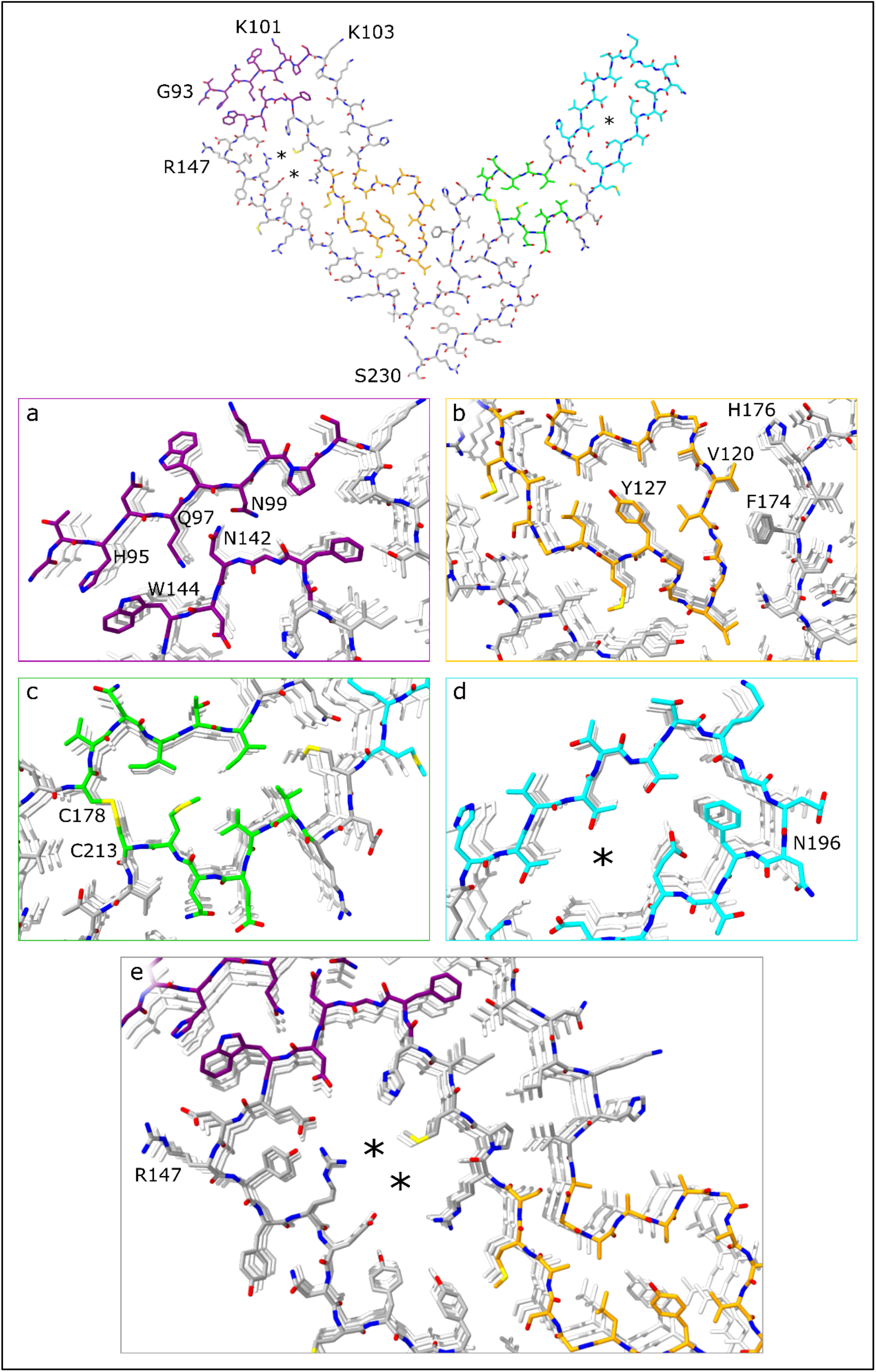
Features of the aRML prion core. **a**, Steric zipper (colored in purple) is formed by the N-terminus and tip of the middle β-arch, inclusive of residues H95, Q97, N99, N142, W144. **b**, a Greek key motif (tan) comprised of hydrophobic residues occurs adjacent to the central strand, stabilized by hydrophobic interface of V120, F174, and H176 residues. **c**, Disulfide bond (C178 – C213) forms the base of disulfide β-arch (green) that is stabilized by a tight interface adjacent to a widening of the arch (**d)**, giving a presumably hydrated pocket (turquoise) marked with an asterisk. **e**, A second hydrated gap occurs in the middle β-arch (grey) of the aRML core, also marked with asterisks.

Like 263K, aRML contains a hydrophobic somewhat Greek key-like motif near the center of the fibril cross-section (Fig. 2a & 3b). However, despite having an identical sequence within residues 112-137 (mouse numbering), the head of the aRML Greek key has a markedly different conformation. A notable example of this is the sidechain of Y127, which is on the inside of the key in aRML, while the analogous Y128 of 263K is on the outside ^2^.

The C-terminal flank of the Greek key arch provides the base of a β-arch of residues ~124-167 (Fig. 2a). The tip of this middle β-arch forms the steric zipper with residues 95-104 described above (Fig. 3a). Although aRML and 263K both have middle β-arches, they differ in sequence at residues 138 and 154 (mouse numbering) and in the conformation of the Greek key residues. Also, the gap between the N- and C-terminal flanks of this arch is wider, and presumably more hydrated, in aRML (Fig. 3e).

A staggered interface between the head of the Greek key and the C-terminal half of the core has the sidechains of residues 120-124 of a given monomer interacting with sidechains of residues 169-176 of the both the same monomer and the one below it. While this stagger is less pronounced than that of 263K ^2^, it contributes to deviations from planarity of each monomer (Fig. 2d).

The β-arch defined by the disulfide bond between C178 and C213 is stabilized by a steric zipper (Fig. 3c). This occurs before a bend and modest widening of the β-arch to encompass a presumably hydrated pocket (Fig. 3d), which is much wider in 263K (Fig. 2c) ^2^.

Another striking difference between the aRML and 263K structures is the orientation of the C-terminal residues that, in 263K, provide a linkage to the GPI anchor. In aRML, the residues 220-230 project away from the disulfide arch and interface with residues 165-170 (Fig. 2 a,b). In contrast, the C-terminal residues of 263K associate with the disulfide arch ^2^.

Comparison of the aRML structure with the 263K prion structure ^2^ reveals that these two *ex vivo* rodent prion strains both have PIRIBS amyloid architectures with several similar structural elements. However, the detailed conformations and relative orientations of these and other elements differ markedly, providing distinct conformational templates for incoming monomers as proposed previously ^2,9^. Also, the amino acid sequences differ at 8 residues within the ordered cores of aRML and 263K ^2^; presumably contributing to the respective templating activities, and, hence, species specificities of these prion strains. In both fibril strains, the opposite ends are not equivalent. Among the key differences is the non-coplanarity of the head of the Greek key β-arch which protrudes at one end and recedes at the other (Fig. 1f & 2d). Such non-equivalence presumably affects the relative mechanism and kinetics of PrP conversion at the fibril ends.

aRML prions lack GPI anchors, but are still lethal ^5^. Thus, neither the GPI anchor, nor the accompanying heavier glycosylation, are prerequisites for neuropathogenicity. From a structural perspective, these post-translational modifications do not seem to alter the core structures of at least 3 murine prions strains, at least as probed by infrared spectroscopy ^3,10–12^. Moreover, *in vivo* passages of RML scrapie from wildtype to anchorless PrP transgenic mice and back again does not alter the strain characteristics of this strain ^5^. Consistent with this conclusion is a structure posted a week ago on a preprint server for wildtype (GPI-anchored and more glycosylated) RML fibrils, which appears to be very similar to that of aRML ^13^. So far, the comparisons of RML prions to 263K prions illustrate the conformational permutations that define two brain-derived prion strains, but much further work is needed to characterize the full spectrum of mammalian prion structures.

## Methods

### PrP^Sc^ fibril purifications

aRML fibrils were purified from brains of transgenic mice expression only GPI-anchorless PrP and characterized as part of previous studies ^3,14^. As detailed in those studies, the mice were housed at RML in an AAALAC-accredited facility. Experiments were in accordance with the NIH RML Animal Care and Use Committee approved protocols (2018-011 or 2016-039).

### Cryo-EM

C-Flat 1.2/1.3 300 mesh copper grids (Protochips, Morrisville, NC) were glow-discharged with a 50:50 oxygen/hydrogen mixture in a Solarus 950 (Gatan, Pleasanton CA) for 10 s. Grids were mounted in a EM GP2 plunge freezer (Leica, Buffalo Grove, IL) and a 3 μl droplet of 0.02% amphipol A8-35 in phosphate buffered saline was added to the carbon surface and hand blotted to leave a very thin film. The tweezers were then raised into the chamber of the plunge freezer, which was set to 22°C and 90% humidity. 3 μl of recently sonicated sample was added to the carbon side of the grid and allowed to sit for 60 s. The sample was subsequently blotted for ~4 s followed by a 3 s drain time before plunge freezing in liquid ethane kept at −180°C. Grids were mounted in AutoGrid assemblies and then loaded into a Titan Krios G3i (Thermo Fisher Scientific, Waltham, MA) with a K3 camera and BioQuantum GIF (Gatan, Pleasanton, CA). Images were acquired at 0.55 Å/pixel at Super Resolution mode, 60 e-/Å^2^, and 60 total frames. Movies were collected using SerialEM ^15^.

### Cryo-electron tomography

C-Flat 1.2/1.3 300 mesh copper grids (Protochips, Morrisville, NC) were glow-discharged with a 50:50 oxygen/hydrogen mixture in a Solarus 950 (Gatan, Pleasanton CA) for 10 sec. Grids were mounted in the tweezers for an EM GP2 plunge freezer (Leica, Buffalo Grove, IL) and a 3 μl droplet of 0.02% Amphipol A8-35 in PBS was added to the carbon surface and hand blotted, leaving behind a very thin film. The tweezers were then raised into the chamber of the plunge freezer, which was set to 22°C and 90% humidity. 3 μl of sample and 1 μl of 5 nm Protein A gold (CMC, Utrecht, The Netherlands) was added to the carbon side of the grid and allowed to sit for 60 seconds. The sample was blotted for ~4 sec followed by a subsequent 3 sec drain time and then plunged into liquid ethane kept at −180°C. Grids were mounted in AutoGrid assemblies and then loaded into a Krios G1 (Thermo Fisher Scientific, Waltham, MA) transmission electron microscope operating at 300 kV with a K3 (Gatan, Pleasanton CA) and a Biocontinuum GIF (Gatan, Pleasanton CA) with a slit width of 20 eV. 13 tilt series were acquired using SerialEM ^15^ at a 0.45 Å pixel size at ±60°, 2° increment in a dose symmetric manner around 0° ^16^ with defocus values ranging from −3 to −6 um and a total dose of ~60 e-/Å^2^. Tomograms were reconstructed and analyzed using IMOD ^17^. The tomographic analyses indicated that all aRML fibrils for which the helical twist was clear (n > 30) were left handed and, as such, a left handed twist was used in the model building described below.

### Image processing of aRML fibrils

Initial map and helical twist parameters were generated based on our recently published dataset ^2^, as well as cryo-electron tomographic analyses of aRML fibrils noted above. Motion correction of raw movie frames was performed with RELION 3.1 ^8^. CTF estimation was performed using CTFIND4.1 ^18^. Fibrils were handpicked then extracted using large and small box sizes. The longer helical segments were extracted with a box size of 1280 pixels and were downscaled to a box size of 256 pixels. The shorter segments were extracted at 400 pixel box size. 2D classes, from the long segments were used to estimate the cross-over distance of the fibril for estimating initial twist parameters. 2D classes from the short segments were used to generate an initial 3D model.

Higher resolution data was collected using a Titan Krios G3i (Thermo Fisher Scientific, Waltham, MA) with a K3 camera and BioQuantum GIF (Gatan, Pleasanton, CA) with images were acquired at 0.55 Å/pixel at Super Resolution mode, 60 e-/Å^2^, and 60 total frames. Movies were motion corrected and the CTF estimated as above. Fibrils were picked manually and segments were extracted with an inter-box distance of 14.6 Å using box size of 740 pixels that was down sampled to 370 pixels. Reference-free 2D class averaging was performed, using a regularization parameter of T = 2, a tube diameter of 180 Å, and the translational offset limited to 4.8 Å. The initial model was used for 3D auto refinement with C1 symmetry, initial resolution limit of 40 Å, initial angular sampling of 3.7°, offset search range of 5 pixels, initial helical twist of −0.72°, initial helical rise of 4.85 Å, and using 50% of the segment central Z length. The output from auto refinement was used for 3D classification without allowing for image alignment to remove poorly aligned segments from auto refinement. Classes were selected for further refinement based on similarity of features in their cross-section (excluding visually low resolution and poorly aligned classes), estimated resolution, overall accuracy of rotation and translation, and Fourier completeness. Auto-refinement was then performed while optimizing the helical twist and rise. Auto-refinement with optimization of twist and rise yielded a final map with a twist of −0.637° and rise of 4.876 Å. Iterative cycles of CTF refinement, Bayesian polishing, and auto refinement were used until resolution estimates stabilized. Post processing in RELION was performed with a soft-edged mask representing 10% of the central Z length of the fibril. Resolution estimates were obtained between independent refined half-maps at 0.143 FSC.

### Model building

De novo building of an aRML atomic model was conducted using Coot ^19^, with the assumption that residues comprising the protease-resistant core (i.e. ~90-231) were included in the amyloid core. Individual subunits were translated to generate a stack of 5 consecutive subunits, and translated subunits rigid-body fit in Coot. Iterative real-space refinement and validation with Coot and Phenix ^20,21^ were carried out, with Fourier space refinements being conducted using RefMac5. Model validation was carried out with CaBLAM ^22^, MolProbity ^23^, and EMringer ^24^, and any outliers/clashes identified and corrected with subsequent iterative refinements/validation. After the addition of protons, the system was solvated in TIP3P water and conjugate gradient energy minimization was performed in NAMD ^25^ utilizing the CHARMM36 forcefield ^26^. A gradient of backbone restraints was utilized in iterative runs of minimization and were subsequently removed in the last 15,000 steps.

## Acknowledgements

We thank Ms. Elizabeth Fisher for helpful suggestions and oversight of the NIH EM facility; Dr. Dave Dorward for help with preliminary negative stain TEM. We thank Dr. Sudha Chakrapani for oversight of the cryo-EM core at the Cleveland Center for Membrane and Structural Biology. We thank Drs. Cathryn Haigh, Parvez Alam, Moses Leavens and Ankit Srivastava for critical review of this manuscript. This work was supported by the Intramural Research Program of the NIAID; Mary Hilderman Smith, Zoë Smith Jaye, and Jenny Smith Unruh in memory of Jeffrey Smith; and the Britton Fund, Case Research Institute, and CWRU School of Medicine. This work utilized the computational resources of the NIH HPC Biowulf cluster (http://hpc.nih.gov).

## Author Contributions

AK and BC supervised the project. All authors designed experiments. GSB purified the PrP^Sc^. AH and AK provided *in vitro* analyzes of PrP^Sc^ preparations. GJR provided brain tissues for PrP^Sc^ purifications. CS and AH prepared EM grids and AK, HS, and KL collected EM data. CS analyzed the tomographic data. FH, CS, BH, and AK processed the cryo-EM images. FH, AK developed map parameters and FH performed class averaging and refined EM map densities in Relion. FH, HS, BC, AK built *de novo* atomic models of PrP^Sc^, FH, HS, and AK performed iterative refinements and map/model validation. EA performed validations for model stability. BC and AK assembled the initial draft and all authors helped with further editing and preparation of the manuscript.

---

**Supplemental Table.**

|  |  |
| --- | --- |
|  | aRML (pdb, emdb) |
| **Data collection and processing** |  |
| Magnification | 81,000x |
| Voltage (kV) | 300 |
| Electron dose (e-/Å<sup>2</sup>) | 60 |
| Pixel size (super resolution) | 0.545 |
| Symmetry imposed | C1 |
| Initial particle segments | 135939 |
| Final particle segments | 15857 |
| Map resolution (Å) (FSC 0.143) | 3.03 |
| Helical rise (Å) | 4.876 |
| Helical twist (°) | -0.637 |
| Map sharpening *B* factor (Å<sup>2</sup>) | 9.28 |
| **Model Refinement** |  |
| Initial Model | *de novo* |
| R.M.S. deviations |  |
| Bond lengths (Å) | 0.002 |
| Bond angles (°) | 0.570 |
| MolProbity score | 1.43 |
| Clash score | 7.94 |
| Rotamer outliers (%) | 0 |
| Ramachandran plot |  |
| Favored (%) | 100.00 |
| Allowed (%) | 0 |
| Outliers (%) | 0 |
| EM Ringer score | 2.90 |
| Model vs. Data (CC) | 0.74 |

**Supplemental Fig. 1.**
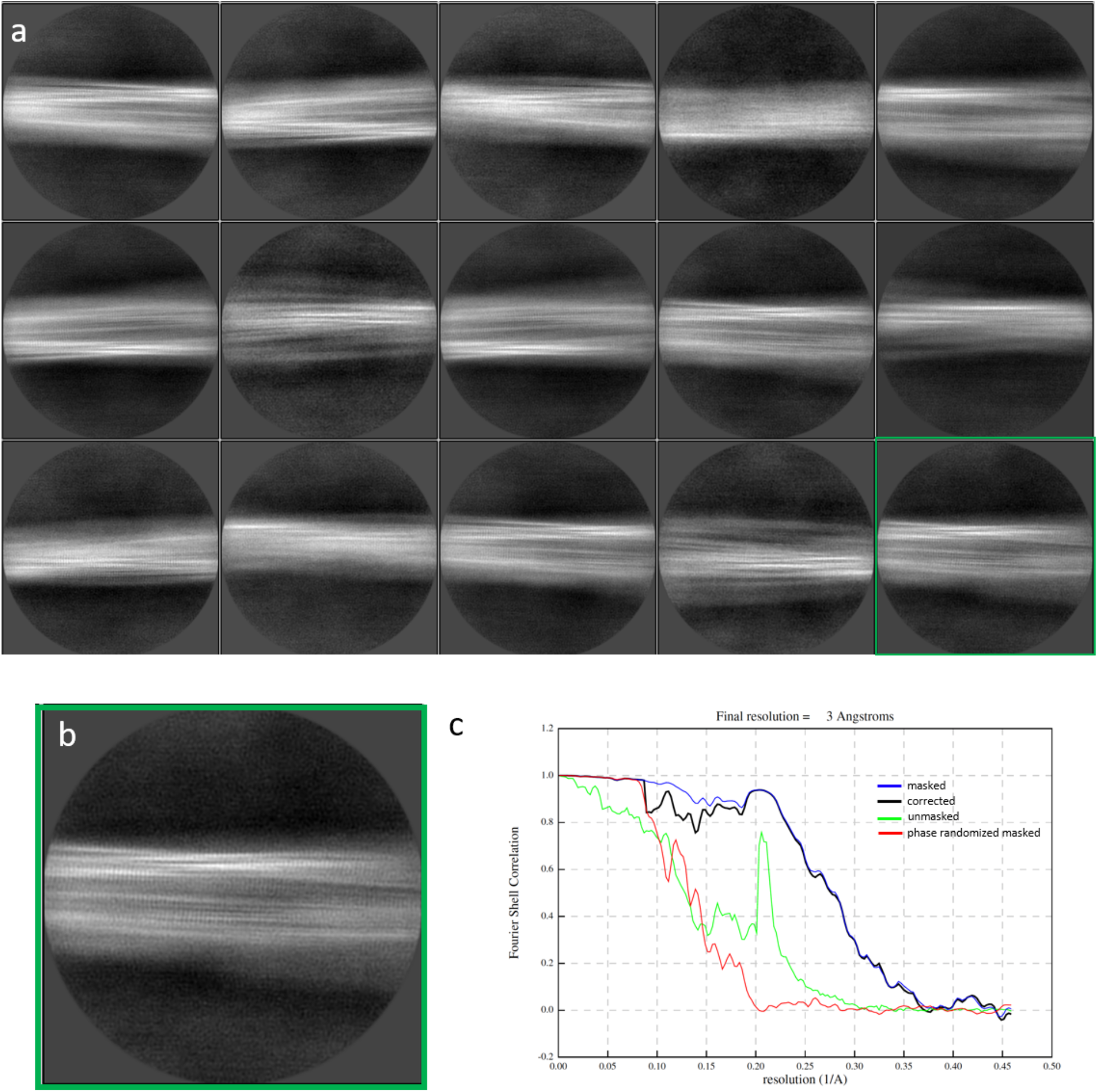
**a**. Representative 2D class averages depicting lateral views of aRML fibril segments (29 classes shown of 50 total used to reconstruct 3D density map). **b**. Enlarged view of one of the 2D classes (highlighted in **a**) showing the 4.9Å repeated spacing perpendicular to the fibril axis. **c**. Fourier shell correlation plots of masked and unmasked models.

**Supplemental Fig. 2.**
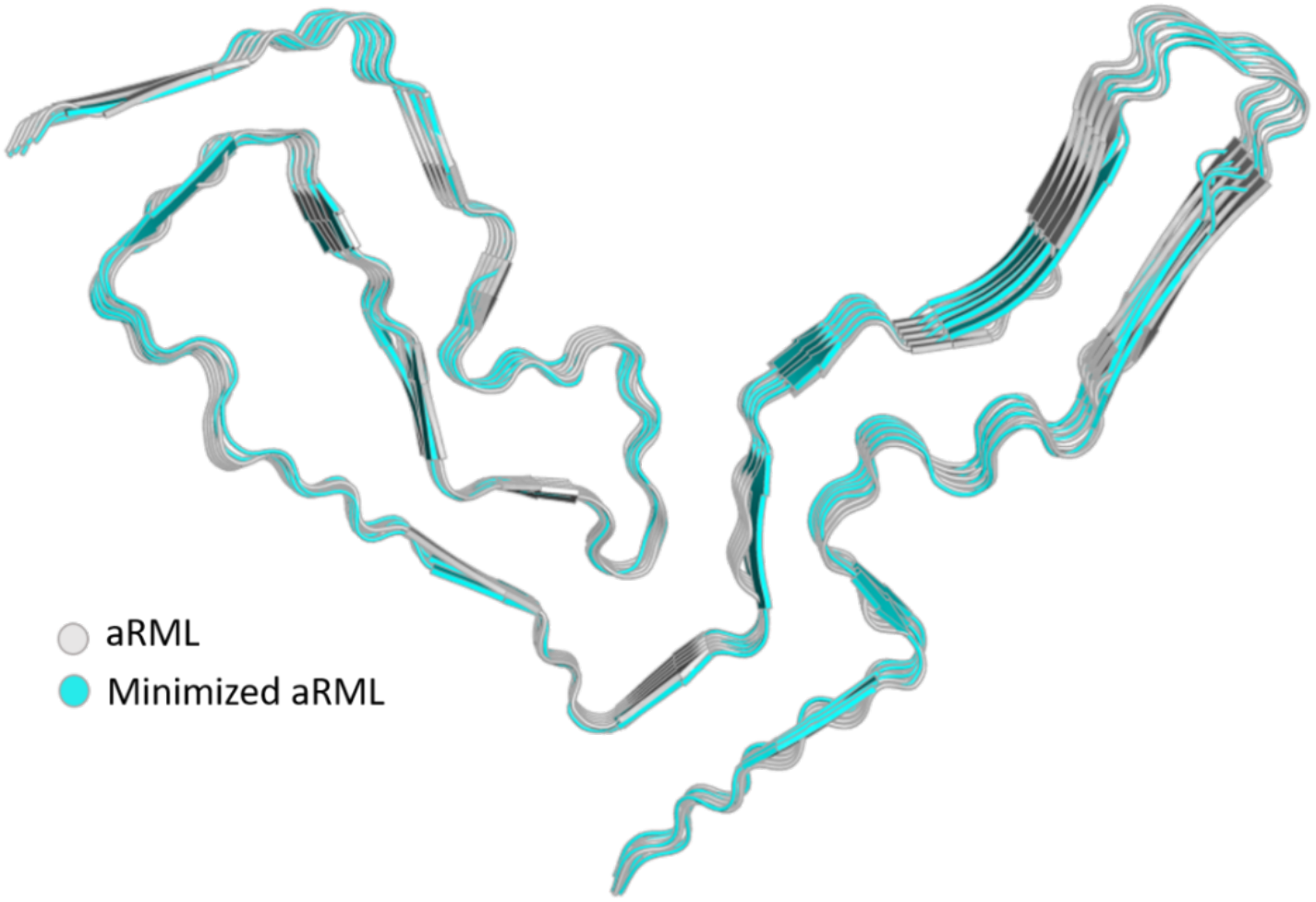
Conjugate gradient energy minimization was utilized to ensure that atomic placement is energetically favorable, and that the conformer is free of steric clashes and other structural anomalies. The relative stability of this structure will be rigorously probed in forthcoming molecular dynamics simulations.

